# *Fusarium graminearum* copper amine-oxidases redundantly increase virulence by converting tryptamine from hydrolyzed plant defense compounds into auxin

**DOI:** 10.1101/2024.10.21.619423

**Authors:** Thomas Svoboda, Asja Ceranic, Pia Spörhase, Anika Bartholomäus, Gerlinde Wiesenberger, Philipp Fruhmann, Eduardo Beltran, Franz Berthiller, Rudolf Krska, Rainer Schuhmacher, Gerhard Adam

**Affiliations:** Institute of Microbial Genetics, Department of Applied Genetics and Cell Biology, University of Natural Resources and Life Sciences, Vienna (BOKU), Konrad-Lorenz-Str. 24, 3430 Tulln, Austria; Institute of Bioanalytics and Agro-Metabolomics, Department of Agrobiotechnology (IFA-Tulln), University of Natural Resources and Life Sciences, Vienna (BOKU), Konrad-Lorenz-Str. 20, 3430 Tulln, Austria

## Abstract

Plant pathogenic fungi have evolved different strategies to interfere with plant defense mechanisms. The well described fungal plant pathogen *Fusarium graminearum* is not only able to produce trichothecene toxins like deoxynivalenol, but also the plant hormone auxin. Highly elevated levels of auxin and auxin derivatives such as IAA-glucoside or IAA amino-acid conjugates were observed in wheat cultivar Apogee infected with *F. graminearum*. We report that *F. graminearum* is able to cleave tryptamine-derived hydroxycinnamic acid amides, e.g. the defense compound coumaroyl-tryptamine. In this study we investigated copper amine-oxidases, candidate genes for auxin biosynthesis converting tryptamine into the IAA precursor indole-3-acetyldehyde. After consecutive knock outs of all seven copper amine oxidases the resulting septuple knock out strain had strongly reduced ability to produce auxin. Virulence of the septuple mutant was significantly impaired while DON production *in planta* was comparable to the wild type. We conclude that *F. graminearum*, often presumed to be a simple nectrotroph, has a biotrophic phase and is able to employ plant defense compounds by converting them into defense suppressing auxin.

## Introduction

Auxin (indole-3-acetic acid, IAA), is a plant hormone involved in several growth related processes as well as abiotic stress response (Gomes and Scortecci 2021; Waadt et al. 2022). The role of auxin in defense responses of plants against microbes is not completely clear yet, and the literature at first glance contradictory (for reviews see (Robert-Seilaniantz, Grant, and Jones 2011; Kunkel and Johnson 2021)). In general, auxin signaling antagonizes salicylic acid (SA) mediated defense signaling (Kunkel and Harper 2018) which is effective against biotrophs and reciprocally, SA inhibits auxin signaling (Du et al. 2013). On the other hand, there is also an antagonism between salicylic acid and jasmonic acid/ethylene signaling, which is effective against necrotrophs. Depending on the plant and pathogen studied different outcomes are observed. Auxin signaling can be necessary for resistance, e.g. impaired auxin signaling in Arabidopsis auxin signaling mutants *axr1, axr2* and *axr6* led to increased susceptibility to the necrotrophic fungi *Plectosphaerella cucumerina* and *Botrytis cinerea* (Llorente et al. 2008). In contrast many microbes are able to synthesize auxin, thereby promoting virulence (Kunkel and Johnson 2021). Auxin may suppress defense also in salicylic acid independent ways (Mutka et al. 2013), for instance in the interaction of *Arabidopsis* with *Fusarium oxysporum* (Kidd et al. 2011).

There is increasing evidence that several fungal effector proteins target the TOPLESS related corepressors for triggering auxin signaling. First described in *Ustilago* (Bindics et al. 2022; Navarrete et al. 2022), this is seemingly also the case for *F. oxysporum* induced wilt diseases in tomato and Arabidopsis, where certain TOPLESS related genes are susceptibility factors (Aalders et al. 2024). Exogenous auxin application has a susceptibility promoting effect upon infection with biotrophic and hemibiotrophic pathogens like *Magnaporthe grisea* (J. Fu et al. 2011) or the oomycete *Phytophhora parasitica* (Evangelisti et al. 2013). Yet, in case of *Fusarium culmorum* and wheat it has been reported that pretreatment with IAA increased resistance of wheat (Petti et al. 2012). Especially in susceptible cultivars, after infection with *F. graminearum* highly elevated levels of auxin were detected in wheat ears (Wang et al. 2018). Using RNAi-mediated knockdown it has been demonstrated that the wheat auxin receptor (TaTIR1) mediates susceptibility (Su et al. 2021) to *F. graminearum*. This study reported also, that application of exogenous auxin increased susceptibility, contradicting Wang et. al (Wang et al. 2018).

Multiple bacteria and fungi exploit the fact that elevated auxin levels increase susceptibility by producing auxin themselves. In this context tryptophan dependent and tryptophan independent pathways have been described (S.-F. Fu et al. 2015). In the indole-3-acetaldoxime (IAOx) pathway, auxin is produced from IAOx which is produced from tryptophan by cytochrome P450 monooxygenases. In the YUCCA pathway, Trp is first converted to indole-3-pyruvic acid (IPA) by an aminotransferase, and then converted into auxin by flavin monooxygenase (encoded by YUCCA genes). In the IAM pathway Trp is first converted to indole-3-acetamide (IAM) which is further metabolized to auxin by indeole-3-acetamide hydrolase (*iaaH, AMI1*). The auxin biosynthetic pathway via i*aaM iaaH* has been reported to be present in certain *Fusarium* isolates in the *Gibberella fujikuroi* species complex (Niehaus et al. 2016; Tsavkelova et al. 2012), but is absent in *F. graminearum*.

The pathway we are focusing on in this study is the one via tryptamine (TAM). In the first step Trp is decarboxylated by a tryptophan decarboxylase to TAM. In the consecutive step TAM is oxidized by an amine oxidase (AOX) resulting in indole-3-acetaldehyde which is further oxidized to auxin (Cao et al. 2019; Mano and Nemoto 2012; Zhao 2010; Sugawara et al. 2009).

Different Fusarium species have been described to be capable of auxin production although different pathways may be used (Tsavkelova et al. 2012; Niehaus et al. 2016). *Fusarium graminearum* itself has been described to be able to produce auxin from TAM and IPA and also to be able to metabolize auxin (K. Luo et al. 2016).

In this study we focus on TAM metabolization by *Fusarium graminearum*. We determine the mechanism of auxin formation, describe the ability of the fungus to cleave hydroxycinnamic acid amides releasing TAM, and assess the impact on disruption on seven copper amine oxidases in *Fusarium graminearum* on auxin production and virulence.

## Results

### *F. graminearum* can cleave CouTam

We tested whether *F. graminearum* PH-1 during infection of the model wheat cultivar Apogee (Mackintosh et al. 2006) increases auxin levels. Upon spray inoculation with *F. graminearum* strain PH-1 after 10 days auxin levels of about 73 μM were measured, compared to about 0.6 μM in the mock treatment. The conjugate of auxin with aspartic acid, IAA-Asp, was elevated from below limit of detection to 62 μM. Likewise, an about 10-fold increase of oxidized auxin (2-oxindole-3-acetic acid) to about 8.5 μM was observed in infected wheat. Likewise, auxin-glucoside (ester), a rather instable substance for which no which no standard is commercially available, was present in high levels while remaining below detection level in the control. In agreement with previous reports (K. Luo et al. 2016) we conclude the *Fusarium graminearum* infection increases auxin levels in wheat ears, and the plant tries to inactivate the excess auxin by glycosylation, oxidation (Hayashi et al. 2021), and formation of IAA-Asp, which is often considered to be an irreversible auxin-inactivation product (P. Luo et al. 2023; Rosquete, Barbez, and Kleine-Vehn 2012).

We next asked what the source for the elevated auxin might be. Already in the first microarray experiment reporting the transcriptome changes in barley after *F. graminearum* infection, it was recognized that the barley tryptophan biosynthesis pathway and also trpytophan decarboxylase were upregulated (Boddu et al. 2006). While (low) increased levels of TAM after Fusarium infection were previously reported for Brachypodium (Pasquet et al. 2014), the main function of TAM seems to serve as precursor for the synthesis of tryptamine derived hydroxycinnamic acid amides (HCAAs) or phenolamids (Liu et al. 2022). In these compounds, a hydroxycinnamic acid (coumaric acid, ferulic acid, caffeic acid … etc.) forms an amide bond with the amino group of tryptamine. Hydroxycinnamic acid amides (HCAAs) are induced upon infection (Kushalappa, Yogendra, and Karre 2016) and presumably have antifungal and defense properties (Doppler et al. 2019; Gunnaiah et al. 2012), yet the experimental evidence is scarce. To test whether *F. graminearum* growth is inhibited we have chemically synthesized coumaroyl-tryptamine (and caffeoyl-tryptamine) and also derivatives with hydroxylated tryptamine (= serotonin), feruloyl-serotonine, coumaroyl-serotoinin using published procedures (Macoy et al. 2022; Takao et al. 2017). An agar diffusion test of *F. graminearum* PH-1 with CouTam is shown in Figure 1. While *F. graminearum* is initially inhibited, the zone of inhibition is decreasing between three and four days of incubation and eventually is overgrown, indicating that the fungus is able to adapt to the plant defense compound and become resistant. In the next experiment, *F. graminearum* growth on minimal media supplemented with different concentrations of CouTam ranging from 0 to 1000 mg/l was tested. The later concentration is already above the limit of solubility and a milky appearance of the agar is observed. It is shown in Figure 1a that the higher the CouTam concentration, the stronger is the delay of germination of conidia. However, once the fungus overcomes initial inhibition it can grow almost without restraint. When the fungus is growing at a high concentration a clearing zone in the milky-appearance ahead of the fungus was observed (Spörhase 2015).

**Figure 1:**
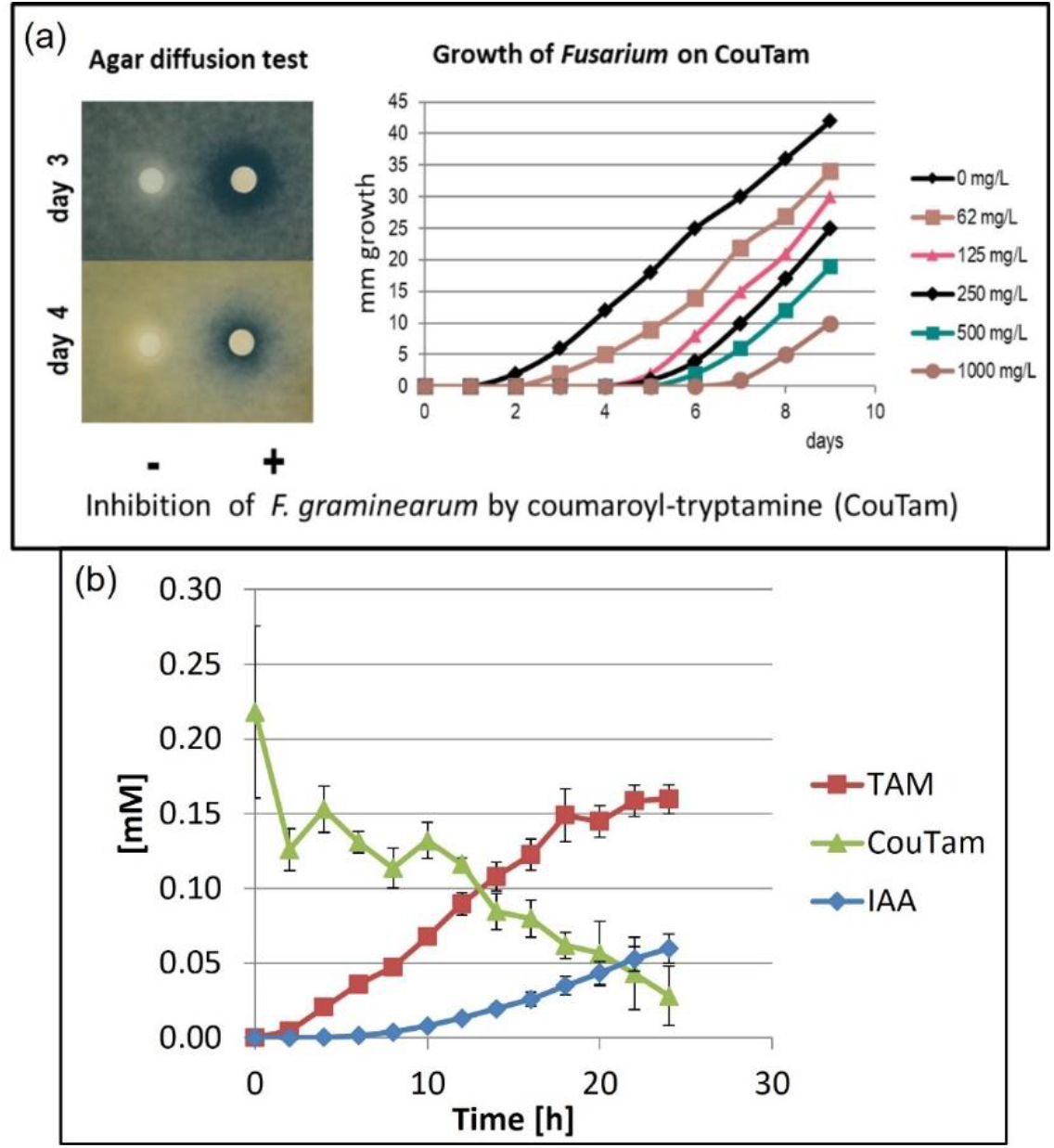
growth inhibition of Fus*arium graminearum* by CouTam (a) and degradation of CouTam over time and formation of auxin (b)

To test whether inactivation of the compound by a putative secreted amidohydrolase cleaving the conjugate is responsible for this phenomenon, a feeding assay in liquid culture was performed. As shown in Figure 1b CouTAM decreased over time while TAM was increasing. We also observed that after eight hours of incubation auxin was formed as well. After 18 hours of incubation TAM hardly increased, while CouTAM was still decreasing and auxin increasing, indicating that the fungus utilizes tryptamine which is released by the cleavage of CouTAM for the formation of auxin.

Once the amidohydrolase is expressed, CouTam is degraded leading to the release of tryptamine. A similar feeding assay with *F. graminearum* and TAM revealed that within 24 hours in liquid culture TAM in nearly 100% yield was converted into auxin. Likewise, when serotonine (5-hydroxy-tryptamine) was fed to it was converted into 5-hydroxy-indole-3-acetic acid (data not shown) which reportedly has much lower auxin activity in oat (Böttger, Engvild, and Soll 1978).

### Inactivation of candidate auxin biosynthetic genes

Assuming that most of the elevated auxin *in planta* is derived from TAM, we selected candidate genes for auxin biosynthesis. In principle amine oxidases are either flavine-dependent enzymes (Pfam: PF01593, EC 1.4.3.4, (Tararina and Allen 2020)) or copper amine oxidases (Pfam01179, EC 1.4.3.21, KEGG reaction R01853). They can remove the amino group from primary amines, generating the corresponding aldehyde, ammonium and hydrogen peroxide: R-CH_2_-CH_2_-NH_2_ + H_2_O + O_2_ = R-CH_2_-CHO + H_2_O_2_ +NH_3_). Based on transcriptome date we selected the copper amine oxidases as candidate genes. Alternatively, auxin biosynthesis theoretically could also be shut down at the consecutive step by disruption of aldehyde dehydrogenases or oxidases where 17 and two genes, respectively, have been annotated (Güldener et al. 2006) . Hence, consecutive knock outs of copper amine oxidases were chosen in the attempt to block fungus-derived auxin biosynthesis. Single knock out strains lacking one amine oxidase were still able to produce auxin, and also double mutants showed only instable reduced ability to produce auxin (data not shown).

In the genome of *F. graminearum* eight amine oxidases have been bioinformatically annotated (**Table 1**). Since one (AOX5) is truncated and nonfunctional, seven remained to be disrupted. RNA-Seq data derived from the susceptible wheat cultivar Remus with the Fhb1 QTL show that amine oxidases are already expressed six hours post infection, while *TRI5* which is responsible for trichothecene production is expressed only 24 hours post inoculation (https://pgsb.helmholtz-muenchen.de/cgi-bin/db2/BOKUnils/index.cgi, accessed 2016-03-20 defunct – Table 1). These data indicate that the fungus attacks the plant on the hormonal level first before trichothecene production is triggered.

**Table 1:** Expression of AOX genes and TRI5 of *Fusarium graminearum* during wheat infection; relative expression levels indicated in FPKM.

| Sequence_ID | Description | Fg_03h | Fg_06h | Fg_12h | Fg_24h | Fg_36h | Fg_48h |
| --- | --- | --- | --- | --- | --- | --- | --- |
| FGSG_15961 | AOX1 | 0 | 33,6 | 66,8 | 298,7 | 197,6 | 135,3 |
| FGSG_03278 | AOX2 | 0 | 0 | 41,7 | 316,7 | 272,7 | 203,1 |
| FGSG_12129 | AOX3 | 0 | 0 | 6,6 | 100,8 | 73 | 56,6 |
| FGSG_02279 | AOX4 | 0 | 0 | 0 | 109,1 | 47 | 38,8 |
| FGSG_01761 | AOX5Ψ | 0 | 0 | 0 | 0 | 0 | 0 |
| FGSG_10587 | AOX6 | 0 | 0 | 0 | 0 | 0 | 54,4 |
| FGSG_10677 | AOX7 | 0 | 0 | 0 | 28,2 | 51,8 | 56,1 |
| FGSG_13604 | AOX8 | 0 | 0 | 0 | 0 | 6,3 | 0,9 |
| FGSG_03537 | TRI5 | 0 | 0 | 0 | 26,7 | 780,2 | 1.001,1 |

For the construction of the multiple knock out strains, a resistance cassette with a positive (hph, nptII or natI) selection genes fused to a negative selection marker (HSVtk) flanked by two loxP sites was used (Twaruschek et al., 2018). HSVtk allows negative selection on media supplemented with 5-fluor-2-deoxyuridine (F2DU) which was used for marker recycling. The flanking regions of the candidate genes were cloned in the respective vector flanking the resistance cassette.

Construction of the septuple knock out strain was started with disruption of AOX3 since it showed the highest activity in native PAGE with tryptamine upon expression in yeast (data not shown). The second gene to be disrupted was AOX4 in the strain already lacking AOX3 followed by cre-recombination for marker recycling and selection on 250 nM 5-fluoro-2-deoxyuridine (F2DU). In the next step, AOX2 was replaced by the resistance cassette natI-HSVtk in the double knock out strain followed by selection on 20 ppm nourseothricin (nat). The fourth gene which was disrupted in the triple knock out strain was AOX6. One PCR confirmed strain which shows a phenotype comparable to the wild type strain was chosen for pop-out of the resistance cassettes that the last three knock outs can be performed in a row. We continued with the fifth knock out where AOX8 was disrupted in the quadruple knock out strain where all resistance cassettes have been popped out. The tryptamine feeding assay which was performed after every single disruption revealed that the quintuple knock-out strain already shows a strongly reduced auxin production compared to the wild type strain. To check whether the amount of auxin can be further reduced by disruption of the remaining two amine oxidase candidate genes, we proceeded with the disruption of AOX7 in the quintuple knock-out strain and finally AOX1 was disrupted.

Figure 2a shows the scheme of the consecutive knock-outs. After disruption of all seven copper amine oxidases, all genes are replaced by a single loxP site. The results of the screening of the AOXΔ^7^ knock out strain with primer pairs flanking the insertion sites are shown in Figure 2b. All PCR fragments show the expected size confirming that all knock-outs and cre-recombination were successful and no chromosomal rearrangements were induced. Some of the wild-type gene fragments with these primers were too big to give amplicons.

**Figure 2:**
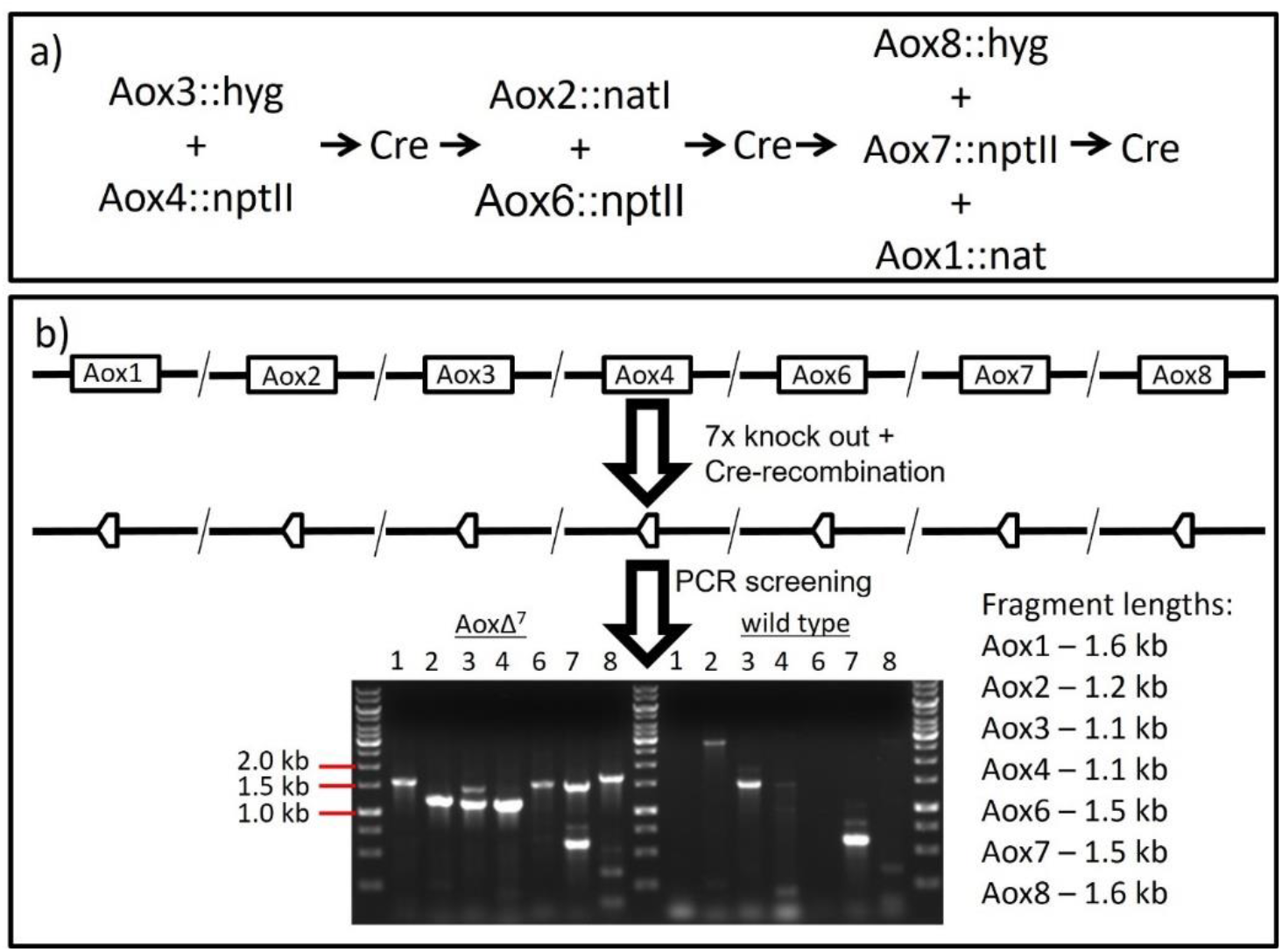
Knock out scheme for the preparation of the septuple knock out strain (a) and PCR confirmation of the septuple knock out (b)

In Supplementary Figure 1 the consecutive knock-outs with the resulting genotypes are shown.

A growth test on potato extract agar (PDA) was performed to determine the growth of the septuple knock out strains taking the wild type into account. As shown in Figure 3f, all tested strains exhibit similar growth as the wild-type.

**Figure 3:**
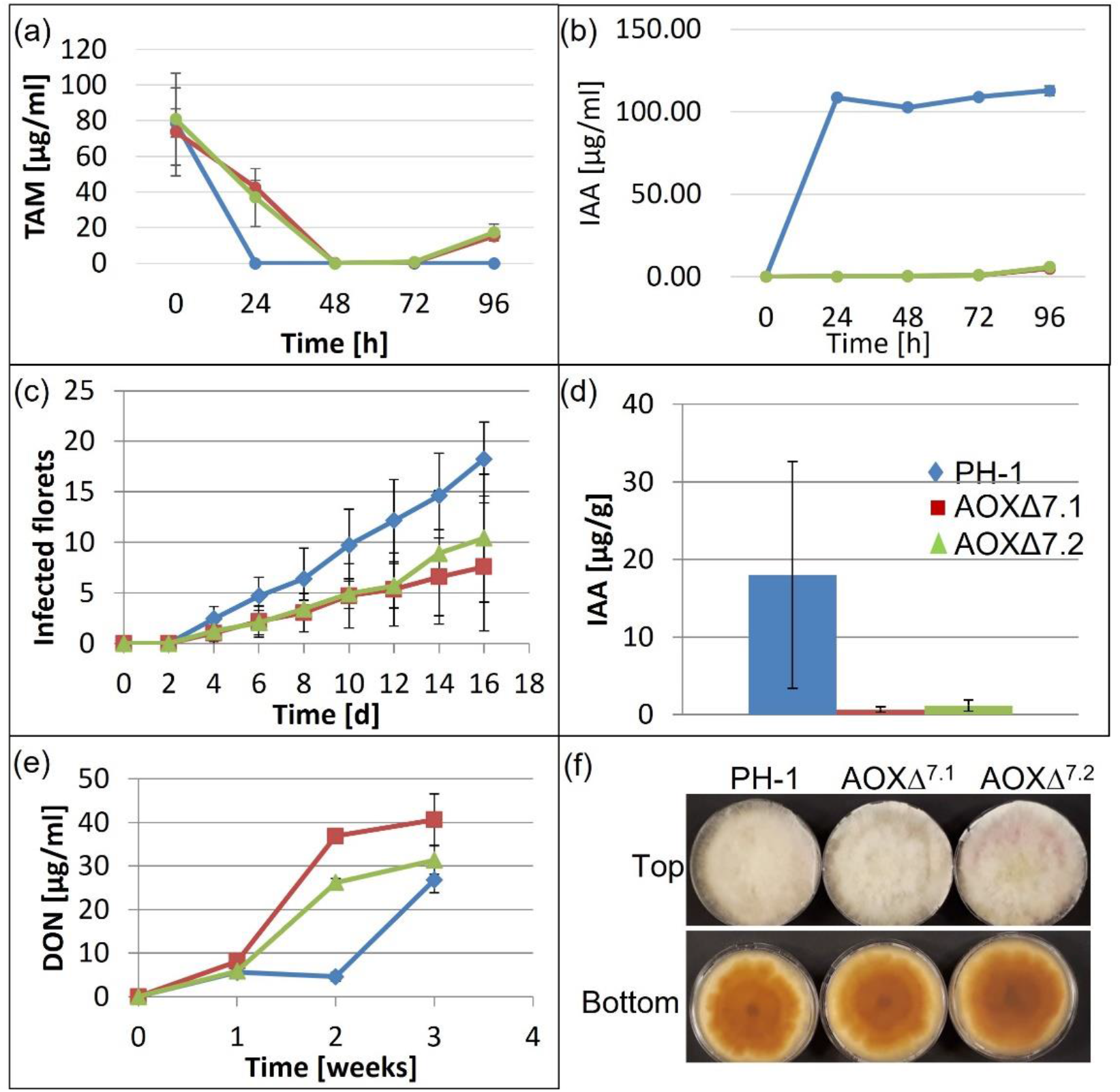
Tests on the septuple knock out strains compared to the wild type strain on TAM consumption (0.5 μM TAM added) (a), and IAA formation in liquid culture (b), progress of infection on the susceptible wheat cultivar Apogee (c), Auxin extracted from infected wheat ears 16 dpi (d), DON formation in liquid culture over time (e), growth on PDA (f)

### TAM metabolization by AOXΔ^7^ strain

Once the septuple knock-out strains were generated, a TAM feeding assay was performed to determine whether the knock-out strains are still able to produce auxin *in vitro*. Two independently derived septuple knock out strains as well as PH-1 wild type as control were inoculated in Fusarium minimal media (FMM) and incubated at 20°C for four days. 1 mM tryptamine was added and samples were taken every 24 hours over a total time period of four days. As expected, TAM was almost completely converted to auxin by the wild type strain within 24 hours. In the cultures containing the AOXΔ^7^ strains, TAM unexpectedly also completely disappeared after 48 hours, however, no auxin was formed at that time. Between 72 and 96 hours about 20% of the initially supplemented tryptamine was released again, indicating that most likely a reversible intermediate was formed, potentially by acetylation or methylation. Low amounts of auxin were detected after 96 hours which is presumably due to the presence of flavin dependent amine oxidases or due to synthesis of auxin via other pathways.

For the virulence tests two florets of 10 ears of the susceptible wheat cultivar Apogee were point inoculated with 10 μl of a 4*10^4^ spores/ml suspension. The progress of infection was documented over a total time period of 16 days. Subsequently, the whole ears were harvested and extracted for determination of the DON and IAA content.

It is shown in Figure 3c that both septuple knock out strains show a significantly slower spread compared to the wild type strain, advancing only throughout less than half the ear, while the wild-type eventually infected all spikelets. Figure 3d shows that the auxin levels in the whole ear are significantly higher in the ears treated with the wild type strain, even though there is a high variation among the wild type treated ears. Interestingly, in the whole ground ear the DON content after 2 weeks was higher, despite lower colonized area than in the wildtype infection, indicating that potentially loss of auxin production may be compensated by higher DON produced *in planta*.

### DON production in liquid medium

To confirm that the lower virulence of the septuple knock out strains is caused by the strongly impaired auxin production, a test on DON production was performed since DON is contributing to virulence as well. The AOXΔ^7^ strains as well as the wild type strain were inoculated in liquid medium suitable for DON production *in vitro*. Therefore, 2SM and incubation for three weeks at 20°C in the dark was done. Samples were taken every week over a total time period of three weeks (Figure 3e). After one week of incubation all strains show comparable DON levels. While the amount of DON in the cultures containing the septuple knock out strains is increasing over time, the DON concentration in the wild type culture is comparable after one week but slightly decreasing after two weeks. After three weeks of incubation the wild type still shows significantly lower DON levels (p<0.05) leading to the conclusion that the DON production is not impaired in the septuple knock out strains.

Aware that the septuple knock out strain produced only low amounts of auxin compared to the wild type strain, we set up an experiment to determine the change of IAA, TAM, TRP and DON levels over time in infected wheat ears. Therefore, two spikelets in the middle of the ear of ten ears per strain and time point were inoculated with 10 ml of a 4*10^4^ spores/ml suspension or 10 μl sterile water as control. After 3, 6, 9 and 12 days, 40 ears, respectively, were harvested and extracted as described in material and methods followed by analysis by LC-MS/MS (Figure 4). As expected, in the ears treated with the wild type strain the auxin levels are significantly higher compared to the ones inoculated with the septuple knock out strains (p<0.05). In contrast, tryptamine levels are significantly lower in the PH-1 wild type treated ears which is in line with our hypothesis since the septuple knock out strains can no longer divert tryptamine into auxin production. After six- and nine-days post inoculation the TAM levels are comparable between all strains indicating that auxin is primarily produced when the fungus penetrates the plant.

**Figure 4:**
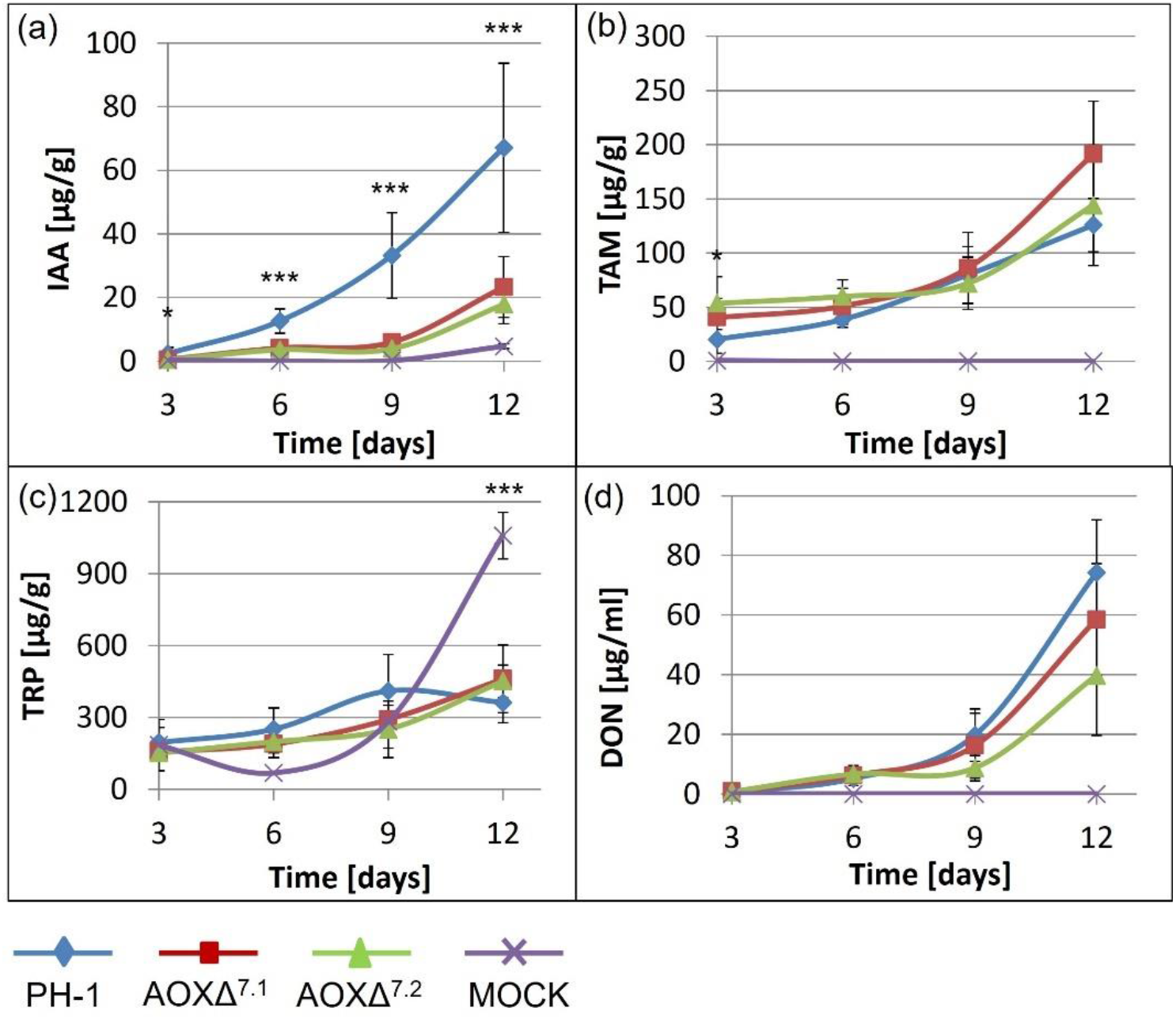
Abundance of auxin (a), TAM (b), TRP (c) and DON (d) over time in infected wheat heads.

The tryptophan levels in the ears inoculated with *Fusarium* are comparable among the strains. Yet, the mock inoculated ears show significantly lower TRP levels 6 dpi, while a strong increase of the tryptophan levels between nine and twelve dpi leads to significantly higher TRP levels compared to the *Fusarium* treated ears. The DON levels are similar until 6 dpi. While AOXΔ^7.1^ shows only slightly lower DON levels compared to the wild type strain, AOXΔ^7.2^ has significantly lower DON levels nine and twelve dpi (p < 0.01). We already showed that the septuple knock out strains show a lower virulence compared to the wild type strain. Since the whole ear was ground and extracted, we can assume that the levels in the infected spikelets are similar or slightly higher than in the wild-type.

### Complementation of AOX4

Since the septuple knock out strain went through multiple transformation procedures, we tested whether auxin biosynthesis can be reconstituted by introducing AOX4. This gene was chosen since it showed the highest enzyme activity with tryptamine when expressed in yeast (Master thesis Pia Spörhase - data not shown). The confirmation of the complementation was done in triplicates using a feeding assay where TAM was supplemented to the liquid media.

The results clearly indicate that the complementation was successful. Both complementation strains metabolize about 100% of the supplemented tryptamine to auxin within one day, like the wild type. The septuple knock out strain forms low amounts of auxin after three days. However, a reversible intermediate must be formed from tryptamine since TAM is close to zero after 2 days of incubation but increases until 4 days that almost 50% of the initially supplemented TAM are released again.

An infection experiment was done to determine whether the virulence of the complementation strains was reconstituted. The infection was observed over a total time period of 12 days since at this time point more than 50% of the inoculated ears were completely infected. After 9 days of incubation one of the complementation strains, Comp. 49, and the wild type show a significantly higher virulence (p<0.05) compared to the septuple knock out strain. After 12 days both complemented septuple knock out strains showed a significantly higher virulence (p<0.05), similar to the wild type strain (p<0.01) compared to the parental septuple knockout strain (Figure 5a).

**Figure 5:**
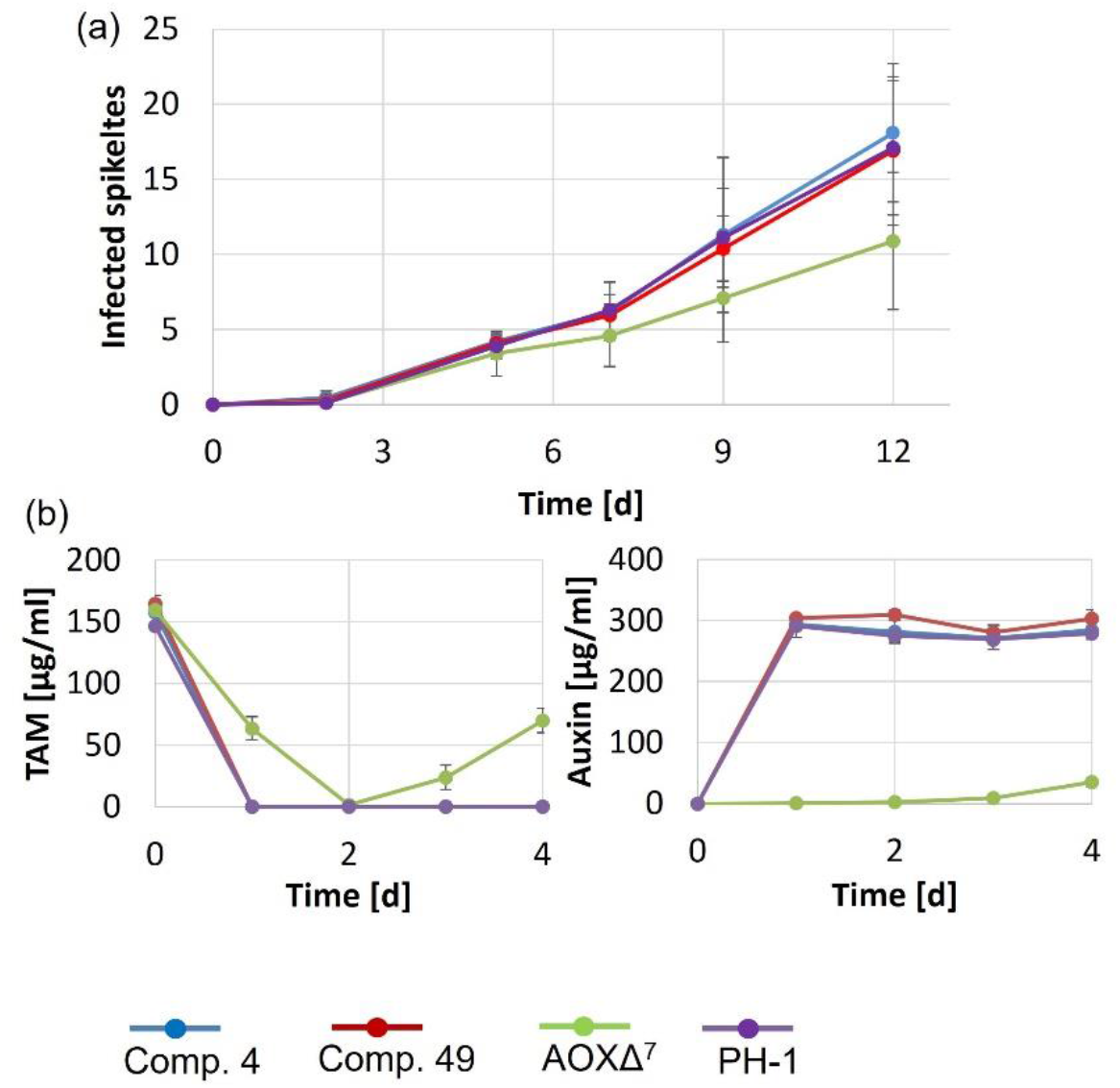
IAA production by complementation strain; wheat infection – fungal spread (a); TAM feeding assay: changes in TAM and auxin levels over time in liquid culture (b)

**Figure 6:**
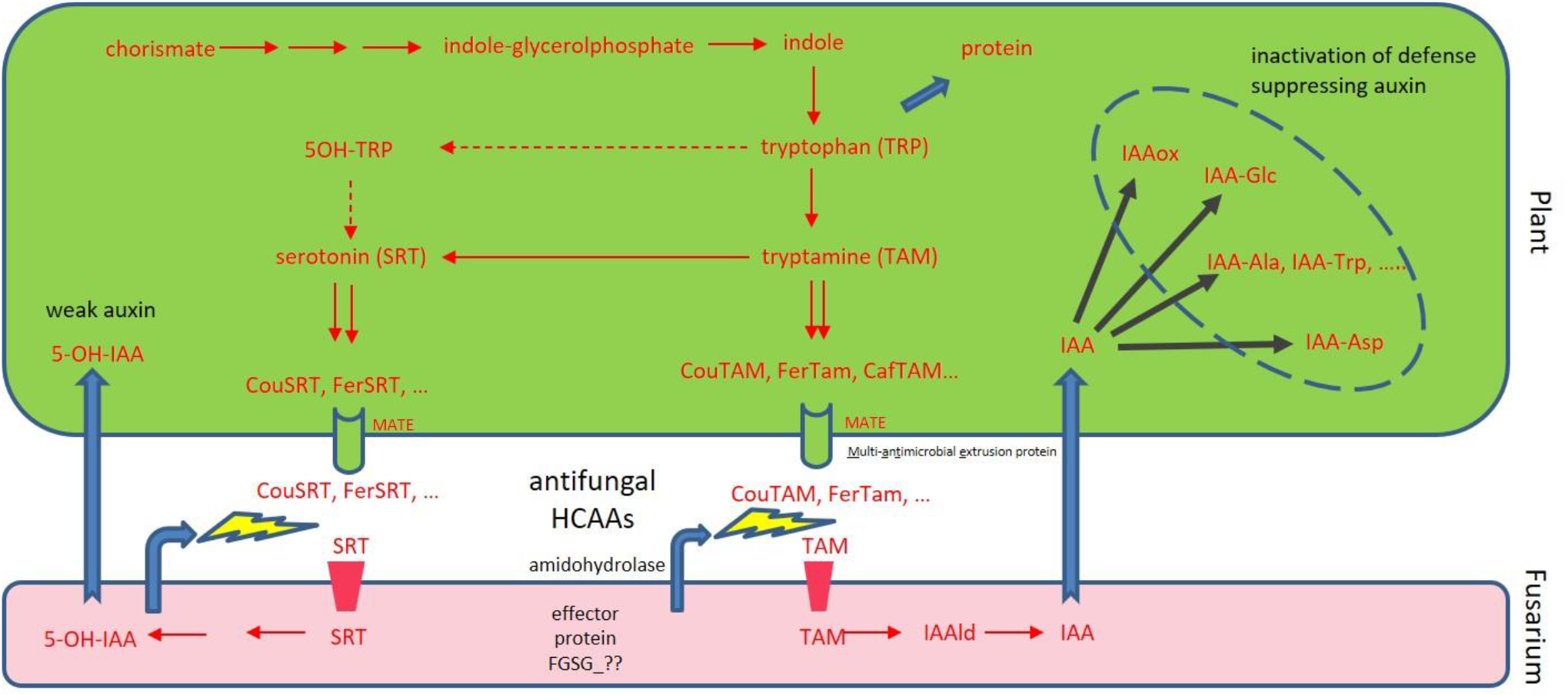
Auxin hypothesis.

## Discussion

*F. graminearum* is often considered to be a necrotroph (Liao et al. 2022), which, in oversimplification, is assumed to produce trichothecene toxins, killing the host plant and then living on the dead tissue. Yet, there is increasing evidence that also *F. graminearum* has an early asymptomatic biotrophic phase (Brown et al. 2017). Our results indicate, that auxin generated early in the interaction is relevant for virulence. We envision a model in which at low fungus density after infection the plants recognize the presence of the invading pathogen, and start to produce hydroxycinnamic acid amides. These compounds have been demonstrated to be antifungal, but also strengthen the cell wall and protect the plant cells against pathogen invasion in this way (Campos et al. 2014). Even though HCAA accumulation induces the hypersensitive response in the plant (Walters 2003) it has been suggested to have an impact on the resistance of wheat towards *Fusarium graminearum* (Gunnaiah et al. 2012). A more recent study suggested that the timepoint of HCAA formation after infection is crucial for a wheat plant determining its resistance level (Whitney et al. 2022). We obtained evidence that coumaroyl-tryptamne (and other HCAAs, synthesized, data not shown) are growth inhibitory. Yet, *F. graminearum* can hydrolyze HCAAs, presumably by secreted amidohydrolases, an area of future research. But it seems reasonable to assume that the direct antifungal activity of the products, tryptamine and courmaric acid, is lower than that of courmaroyl-tryptamine and this is already a detoxification process. Secondarily. the tryptamine released is a convenient nitrogen source if taken up and metabolized by peroxisomal amine-oxidases. *F. graminearum* can use it as sole carbon and nitrogen source (Spörhase 2015), but TAM also gets toxic at high concentrations above 125 mg/L (about 0.8 mM), which is in the highest range observed *in planta* (Fig 4b). Yet, the fungus is able to convert TAM into the plant growth hormone IAA, that at much lower concentrations is hormonally active and suppresses defense. IAA has been reported to be growth inhibitory for *F. graminearum* in high concentrations (0.1-1 mM (∼175 mg/L)) *in vitro* (K. Luo et al. 2016), and auxin can also be metabolized. In the infection experiment shown in Figure 4 maximum levels above 60 mg/L IAA have been detected, so the range where the fungus is also negatively affected can be reached. The high auxin wave should diffuse faster than the fungal infection front moves, thereby conditioning a state of susceptibility. The high auxin triggers its inactivation by induction of transcripts encoding enzymes involved in glycosylation, amino-acid conjugation and oxidation. Yet, the production of active enzymes could be blocked or delayed by the trichothecene toxin DON, blocking protein synthesis.

The main problem with Fusarium infection is the contamination of grain with the trichothecene mycotoxin deoxynivalenol. One would expect that DON-levels in ears infected by the septuple mutant, should be about half if only half of the ear is colonized, which was not observed. TAM, in contrast to other amines, that are converted into HCAAs, like putrescine and agmatine, does only have (if at all) a small stimulatory effect in DON production *in vitro* (K. Luo et al. 2016; Gardiner et al. 2010). Yet, by knocking out all amine oxidases, theoretically DON-inducing other amines may also accumulate to higher levels, which could explain the unchanged or even higher DON levels observed, despite a smaller portion of the ear was colonized. This phenomenon should to be addressed in future research.

The loss of the amine oxidases leads to lower virulence, indicating that auxin suppresses defense, supposedly by antagonizing SA-mediated defense (Hao et al. 2018; Makandar et al. 2012), which is also targeted directly by fungal effectors capable to inactivate SA (Qi et al. 2019). A relevant question is how widespread the hijacking of auxin signaling via amine-oxidase mediated auxin production from TAM is in fungal pathogens. Amine-oxidases are present in many fungi, and the mechanism described her seems to be widespread and redundant. We have observed that for instance many members of the *Fusarium oxysporum* and *Gibberella fujikuroi* species complex *in vitro* can efficiently produce auxin in tryptamine supplemented medium. *In planta*, the limiting factor is most likely the ability to hydrolyze the HCAAs.

In the septuple aminoxidase deficient mutant still residual auxin production after extended time *in vitro* was observed, and also *in vivo* lower, but still elevated levels compared to mock treated plants were observed (Fig 4a). Potentially also flavine-dependent amine oxidases might contribute to TAM-dependent auxin synthesis. Yet, most likely, the fungus can also tap into the high plant tryptophan pool (Fig 4C) and synthesize auxin in other ways – candidate Fusarium YUCCA-like genes exist, that are still uncharacterized.

Wheat plants upon Fusarium infection do not only accumulate TAM-containing HCAAs, but also high levels of serotonin-derived metabolites. While serotonin is derived from hydroxylated-tryptamine in animals, TRP is first decarboxylated and then tryptamine can be hydroxylated to generate serotonin in plants. When compounds like feruloyl-serotonin are hydrolyzed, the released serotonin can also be a substrate of amine-oxidases and via aldehyde-dehydrogenase, 5-Hydoxy-IAA is generated in high yield. Yet, when this metabolite is turned against the plant, it has much weaker auxin activity. In oat it has been reported that 5-HIAA has about 1% of the activity of DON. So, a hypothesis to be tested in future work is that plants which have higher ability to shift TAM into SER and the corresponding HCAAs have higher Fusarium resistance. Likewise of importance is the ability of plants to cope with the highly elevated auxin levels triggered by the fungus. Processes like glycosylation, oxidation and amino-acid conjugation of auxin my therefor be relevant for resistance breeding. In so far unsuccessful attempts to identify the relevant amidohydrolases, we found that one Fusarium effector protein has the ability to hydrolyze the IAA-Asp conjugate (Manuscript in preparation). Seemingly auxin inactivation is an important battleground in the interaction of the pathogen and the host. Another possibility of plants to counteract auxin is the ability to desensitize auxin signaling components (e.g. the main auxin receptor TIR, (Su et al. 2021) by RNA-interference with microRNAs (P. Luo et al. 2022). Yet, for plant breeders it is no option to trade Fusarium resistance to altered grain development and yield losses. Auxin is highly relevant for developmental processes leading to embryo development and grain filling (Song et al. 2024). To find the right balance between growth and resistance remains a difficult task (Denancé et al. 2013), and the insights from this work can hopefully aid the goal to reduce mycotoxin contamination of grain.

## Conclusion

We showed that *Fusarium graminearum* PH-1 is able to cleave Cou-TAM, an HCAA which is formed by plants upon infection as defense response. Furthermore, we demonstrated that the released TAM is metabolized to auxin. We assessed the auxin, TAM, Trp and DON levels in wheat ears infected with the wild type and both septuple knock out strains. Our results lead to the conclusion that on one side auxin produced by the pathogen is indeed promoting virulence but also that Fusarium is able to cleave the plant defense compound Cou-TAM and use the released TAM as substrate for auxin production. This indicates that the pathogen is not only interfering with plant defense upon infection but also degrading defense compounds of the plant, leading to imbalanced hormone homeostasis and susceptibility.

## Material and Methods

### Growth test on CouTam

For the growth test, spores were generated by inoculation of mycelium scratched from a plate in 50 ml Mung Bean Soup (MBS). The samples were incubated at 20°C, 140 rpm for three days. For the agar diffusion test, sterilized filter papers were soaked with 10 μl of a 0.5 mM CouTAM solution (2% DMSO as control). For the growth test, FMM plates containing different amounts of CouTAM ranging from 0 mg/l to 1000 mg/l were prepared and the diameter of the mycelium was measured every day over one week.

### Plasmid preparation for the knock-outs

The flanking regions of the respective gene were amplified by PCR followed by cloning in the plasmids as shown in Table 1.

**Table 1:** Cloning strategy of the flanking regions of AOX candidate genes. The vectors used in this experiment have been described by Twaruschek et al., 2018.

| Gene | Name | Region | Cloning sites | Primer pairs | Vector |
| --- | --- | --- | --- | --- | --- |
| FGSG_15961 | AOX1 | 5′ | BcuI/SfiI | 2581 + 2582 | pKT247 |
|  |  | 3′ | HindIII/SalI | 2583 + 2584 |  |
| FGSG_03278 | AOX2 | 5′ | BcuI/SfiI | 2827 + 2828 | pKT247 |
|  |  | 3′ | HindIII/SalI | 2829 + 2830 |  |
| FGSG_12129 | AOX3 | 5′ | BcuI/SfiI | 2779 + 2780 | pASB43 |
|  |  | 3′ | HindIII/SalI | 2781 + 2782 |  |
| FGSG_02279 | AOX4 | 5′ | BcuI/SfiI | 2783 + 2784 | pKT245 |
|  |  | 3′ | SplI/SalI | 2785 + 2786 |  |
| FGSG_10587 | AOX6 | 5′ | BcuI/SfiI | 3003 + 3004 | pKT245 |
|  |  | 3′ | HindIII/SalI | 3005 + 3006 |  |
| FGSG_10677 | AOX7 | 5′ | BcuI/SfiI | 3089 + 3090 | pKT245 |
|  |  | 3′ | HindIII/SalI | 3091 + 3092 |  |
| FGSG_13604 | AOX8 | 5' | BcuI/SfiI | 2831 + 2832 | pKT248 |
|  |  | 3' | HindIII/SalI | 2833 + 2834 |  |

The transformation and marker recycling were performed as described by Twaruschek et al. (2018).

### IAA formation – time course

To test the knock out strains on their ability to produce auxin, a pre-culture was prepared by inoculating 1*10^5^ spores of PH-1 wild type and PCR confirmed single and multiple knock out strains in 50 ml Fusarium minimal media (FMM) which was incubated at 20°C, 180 rpm for 3 days. 1 mM tryptamine (TAM) was added to each culture and samples were taken at time point 0 and then every 24 hours over four days. The experiment was carried out in three replicates.

For sampling 1 ml of the culture was transferred to a 1.5 ml tube and centrifuged for 1 minute at full speed. 500 μl of the supernatant were transferred to another tube containing 500 μl methanol. The samples were vortexed for 30 seconds and then centrifuged at full speed for five minutes. 50 μl of these samples were transferred to a HPLC vial containing 950 μl methanol resulting in a final dilution of 1:40 for the measurements. The samples were measured by LC-MS.

### Virulence tests

For the virulence tests the susceptible wheat cultivar Apogee was used. The strains to be tested were sporulated in 50 ml MBS followed by incubation at 20°C, 140 rpm for three days. Subsequently, the spores were filtered, counted and diluted to a final concentration of 4*10^4^ spores/ml. Four florets of two spikelets in the middle of the ear were inoculated with 10 μl of the spore suspension. The ears were wrapped with a plastic bag which was previously sprayed with water to provide humidity and incubated at 20°C with a day/night cycle 16h/8h. The progress of infection is recorded every two days.

### AOX4 complementation

For the complementation, AOX4 was amplified using primer 5’-CTTTGACAATGCTGGTAGATG -3’ and 5’-ATTGTTATCACATATCGCATTAC -3’ with an annealing temperature of 55°C and an extension time of 128 seconds. The final plasmid consists of AOX4 flanked by the 5’- and the 3’-flanking region with nptII adjacent. As prokaryotic selection marker ampicillin was used. Prior transformation of the *Fusarium graminearum* Δ^7^ strain, the plasmid was linearized using SalI. Two independent complementation transformants were obtained. The integration of AOX4 was determined by a TAM feeding assay in liquid culture. Therefore, 1*10^5^ spores of selected transformants as well as the septuple knock out- and the wild type strain were inoculated in 20 ml FMM. After three days of incubation at 20°C, 180 rpm 0.5 mM Tam was supplemented. Samples were taken every 24 hours over a total time period of 4 days starting with 0 hours. For the measurements 1:40 dilutions were prepared.

