## Supplementary figures and images for "*Fusarium graminearum* copper amine-oxidases redundantly increase virulence by converting tryptamine from hydrolyzed plant defense compounds into auxin"

### Supplementary Figure 1

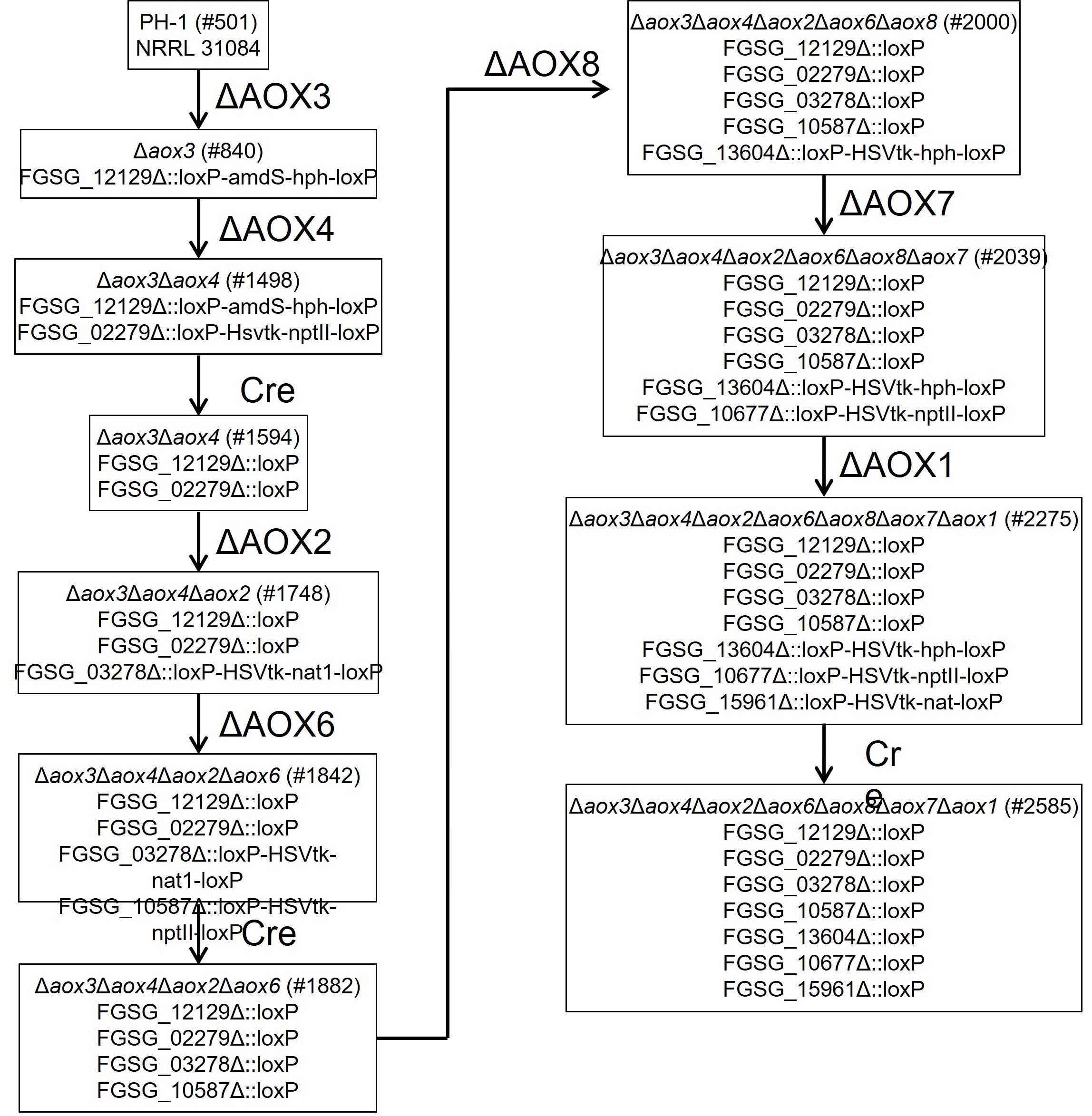
